# Tropomyosin and α-actinin cooperation inhibits fimbrin association with actin filament networks in fission yeast

**DOI:** 10.1101/169961

**Authors:** Jenna R. Christensen, Kaitlin E. Homa, Meghan E. O’Connell, David R. Kovar

## Abstract

We previously discovered that competition between fission yeast actin binding proteins (ABPs) for association with F-actin helps facilitate their sorting to different F-actin networks. Specifically, competition between actin patch ABPs fimbrin Fim1 and cofilin Adf1 enhances each other’s activities, and rapidly displaces tropomyosin Cdc8 from the F-actin network. However, these interactions don’t explain how Fim1, a robust competitor, is prevented from associating equally well with other F-actin networks. Here, with a combination of fission yeast genetics, live cell fluorescent imaging, and in vitro TIRF microscopy, we identified the contractile ring ABP α-actinin Ain1 as a key sorting factor. Fim1 competes with Ain1 for association with F-actin, which is dependent upon their residence time on F-actin. Remarkably, although Fim1 outcompetes both contractile ring ABPs Ain1 and Cdc8 individually, Cdc8 enhances the bundling activity of Ain1 10-fold, allowing the combination of Ain1 and Cdc8 to inhibit Fim1 association with contractile ring F-actin.

## INTRODUCTION

Like other cell types, the unicellular fission yeast assembles diverse actin filament (F-actin) networks within a crowded common cytoplasm to facilitate different cellular functions such as cytokinesis (contractile ring), endocytosis (actin patches) and polarization (actin cables). Each of these F-actin networks possesses a unique overlapping set of actin binding proteins (ABPs) that regulate the network’s formation, organization, and dynamics. However, the mechanisms by which different sets of ABPs sort to particular F-actin networks are less clear. We hypothesize that a combination of competitive and cooperative interactions between different ABPs for association with F-actin may be a driving force in establishing and maintaining their sorting. We previously identified competitive binding interactions between three ABPs with distinct network localizations—fimbrin Fim1 and ADF/cofilin ADF1 (endocytic actin patches) and tropomyosin Cdc8 (cytokinesis contractile ring) (hereafter called Fim1, Adf1 and Cdc8)— that help facilitate their sorting to the proper F-actin networks in fission yeast (Skau and Kovar, 2010; Christensen *et al*., 2017). Specifically, we discovered that synergistic activities between Fim1 and Adf1 rapidly displace Cdc8 from F-actin networks such as endocytic actin patches (Skau and Kovar, 2010; Christensen *et al*., 2017). Although Fim1 prevents Cdc8 from associating with endocytic actin patches, it is unclear how Fim1 is prevented from associating with other F-actin networks such as the contractile ring. Therefore, we sought to determine whether other ABPs at the contractile ring prevent Fim1 association. In this study, we demonstrate that Fim1 competes with the contractile ring ABP α-actinin Ain1 (hereafter called Ain1) for association with F-actin, and that their ability to compete is defined by their residence time on F-actin. Additionally, we show that although Fim1 outcompetes both Cdc8 and Ain1 individually, Cdc8 enhances Ain1-mediated F-actin bundling ten-fold, allowing the combination of Cdc8 and Ain1 to compete with Fim1 for association with F-actin.

## RESULTS

### F-actin crosslinking proteins Fimbrin Fim1 and α-actinin Ain1 compete at the contractile ring and at actin patches

We speculated that competition with contractile ring ABPs prevents Fim1 from strongly associating with the contractile ring. Increasing the percentage of soluble Fim1 might therefore allow Fim1 to outcompete its contractile ring competitors. As Fim1 is concentrated in actin patches (Nakano *et al*., 2001; Wu *et al*., 2001), actin patch depletion by treatment with the Arp2/3 complex inhibitor CK-666 (Nolen *et al*., 2009; Burke *et al*., 2014) is expected to result in a rapid increase of free Fim1 in the cytoplasm. When fission yeast cells expressing Lifeact-GFP were treated with CK-666, we observed a depletion of actin patches and the formation of excessive ‘ectopic’ actin cable and contractile ring material (Figure 1A) (Burke *et al*., 2014).

**Figure 1.**
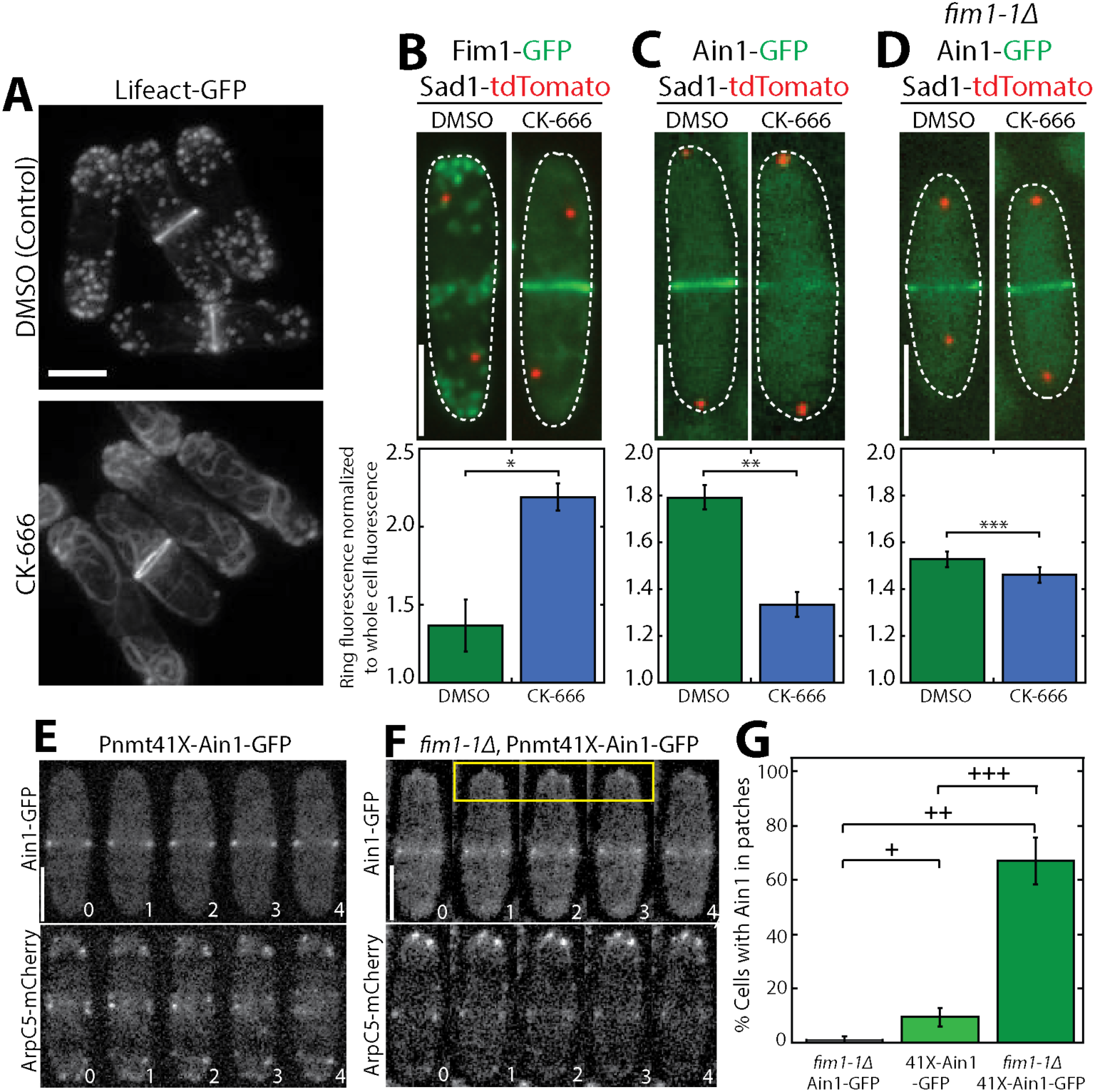
Fimbrin Fim1 and α-actinin Ain1 compete for association with the contractile ring and actin patches. **(A)** Fluorescence micrographs of fission yeast cells expressing Lifeact-GFP following treatment with DMSO (control, top) or 200 μM Arp2/3 complex inhibitor CK-666 (bottom). **(B-D, top panels)** Fluorescence micrographs of fission yeast cells expressing Fim1-GFP **(B)**, Ain1-GFP **(C)**, or Ain1-GFP in a *fim1*-*1Δ* background **(D)**, following treatment with DMSO (left) or 200 μM CK-666 (right). Dotted lines outline cells. Scale bars, 5 μm. **(B-D, bottom panels)** Mean Fim1-GFP **(B)**or Ain1-GFP **(C,D)** fluorescence at the contractile ring normalized to whole cell fluorescence. Error bars=s.e. Two-tailed t-tests for data sets with unequal variance yielded p-values ^∗^p=2.35×10^−5^, ^∗∗^p=1.11×10^−5^, ^∗∗∗^p=0.017. n≥13 cells total in each condition from two independent experiments. **(E-F)** Time-lapse fluorescent micrographs of fission yeast cells expressing ArpC5-mCherry (bottom) and overexpressing GFP-tagged α-actinin Ain1 from the 41xnmt promoter (top) for 20 hours in a wild-type **(E)** or *fim1*-*1Δ* background **(F).** Yellow box highlights Ain1-GFP localization at actin patches. Scale bars, 5 μm. Time in sec. **(G)** Percentage of cells in which Ain1-GFP is observed in actin patches. Error bars=s.e. Two-tailed t-tests for data sets with unequal variance yielded p-values ^+^p=0.113, ^++^p=0.002, ^+++^p=0.012. n=3 experimental replicates.

In control (DMSO-treated) cells, Fim1-GFP localized predominantly to actin patches, with only a small amount associating with the contractile ring (Figure 1B, left) (Wu *et al*., 2001). However, following CK-666 treatment, Fim1-GFP strongly associated with the contractile ring and to a subset of ectopic F-actin (Figure 1B, right). The localization of most contractile ring ABPs, including formin Cdc12, type II myosin Myo2, myosin regulatory light chain Rlc1, the IQGAP Rng2 and tropomyosin Cdc8, was unaffected by CK-666 treatment, (Figure 1–figure supplement 1 and Figure 1–figure supplement 2). Conversely, Ain1 was depleted from the contractile ring following CK-666 treatment (Figure 1C). Therefore, we hypothesized that Fim1 and Ain1 are competitors, and that enhanced Fim1 association with the contractile ring following CK-666 treatment displaces Ain1. We tested this hypothesis by observing Ain1 localization in a strain lacking Fim1 (*fim1*-*1Δ*, Ain1-GFP). In the absence of Fim1, Ain1-GFP was not displaced from the contractile ring following CK-666 treatment (Figure 1D). Localization of Fim1 to the contractile ring and displacement of Ain1 from the contractile ring occurred at all stages of contractile ring assembly and constriction (Figure 1-figure supplement 3).

If competition between Fim1 and Ain1 is a primary driver of their sorting to distinct F-actin networks, we expected to observe Ain1-GFP erroneously localized at actin patches in the absence of Fim1. However, we observed Ain1-GFP at actin patches in less than 1% of *fim1*-*1Δ* cells (Figure 1G, Video 1). It is possible that a combination of the low number of Ain1 molecules (~3,600±500, (Wu and Pollard, 2005)), and the high density of F-actin in actin patches (5,000-7,000 actin molecules in each of 30-50 actin patches, (Wu and Pollard, 2005; Sirotkin *et al*., 2010)), may dilute the Ain1-GFP signal beyond detection. Therefore, increasing the concentration of Ain1-GFP might allow observable Ain1-GFP at actin patches, but only in a *fim1*-*1Δ* background. Indeed, Ain1-GFP overexpressed under the 41xnmt promoter was observed to localize to actin patches in ~67% of *fim1*-*1Δ* cells (Figure 1F-G, Video 1), but in only ~10% of cells expressing endogenous Fim1 (Figure 1E,G, Video 1). These findings indicate that Ain1 is less able to associate with actin patches in the presence of Fim1, suggesting that Fim1 and Ain1 compete for association with F-actin in actin patches as well as the contractile ring.

### Fimbrin Fim1 and α-actinin Ain1 dynamics on F-actin define their competitive effectiveness

Ultrastructural and mutational studies of fimbrin/plastin and α-actinin from several organisms revealed that they bind to a similar site on F-actin (Holtzman *et al*., 1994; Honts *et al*., 1994; McGough *et al*., 1994; Galkin *et al*., 2010; Galkin *et al*., 2008). However, while fission yeast fimbrin Fim1 is relatively stable on single filaments (k_off_= 0.043±0.001 s^−1^) and very stable on F-actin bundles (k_off_=0.023±0.003 s^−1^) (Skau *et al*., 2011), α-actinin Ain1 has not been observed on single filaments and is extremely dynamic on F-actin bundles (k_off_ =3.33 s^−1^ on two-filament and three-filament bundles) (Li *et al*., 2016). Therefore, we hypothesized that Fim1’s longer residence time on F-actin bundles may explain its ability to outcompete Ain1 for the same F-actin binding site. To test this possibility we took advantage of the Ain1 mutant Ain1(R216E), which is less dynamic on F-actin bundles (k_off_ =0.67 s^−1^ and k_off_ =0.33 s^−1^ on two- and three-filament bundles, respectively) (Li *et al*., 2016), and assessed its ability to compete with Fim1 in vitro and in vivo.

We utilized multi-color TIRF microscopy (TIRFM) to visualize fluorescently labeled Fim1 in the presence or absence of unlabeled Ain1 on actin filaments. 50 nM Fim1-TMR fully decorated actin bundles and was also present on single actin filaments (Figure 2A-B). Compared to Fim1-TMR alone, less Fim1-TMR was associated with single actin filaments and two filament F-actin bundles in the presence of either 1 μM wild-type Ain1 or Ain1(R216E) (Figure 2A-B). However, there was little difference in the amount of Fim1-TMR associated with single filaments or two-filament bundles in the presence of wild-type Ain1 or mutant Ain1(R216E).

**Figure 2:**
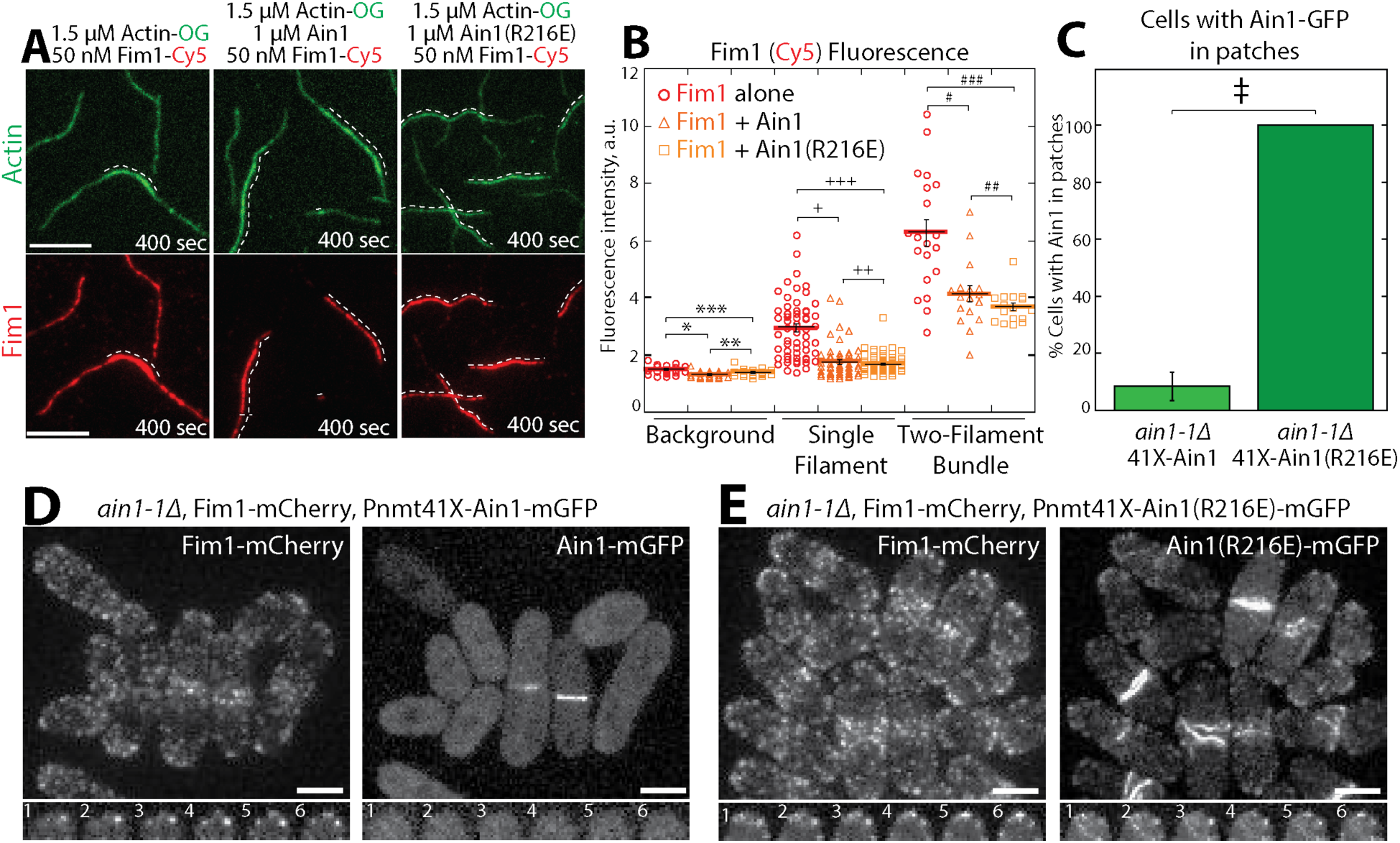
Fimbrin Fim1 and α-actinin Ain1 competition is driven by their residence time on F-actin. **(A-C)** Two-color TIRFM of 1.5 μM Mg-ATP actin (15% Alexa 488-labeled) with 50 nM fimbrin Fim1 (Cy5-labeled) alone, or with either 1 μM wild-type α-actinin Ain1 or mutant Ain1(R216E). Scale bars, 5 μm. Dotted lines denote bundled regions. **(B)** Dot plots of the amount of Fim1-Cy5 fluorescence on the background (coverglass), single actin filaments, or two-filament F-actin bundles in either the absence (red circles) or presence of Ain1 (orange triangles) or Ain1(R216E) (yellow squares). Error bars=s.e. Two-tailed t-tests for data sets with unequal variance yielded p-values ^∗^p=2.16×10^−4^, ^∗∗^p=0.054, ^∗∗∗^p=0.026, ^+^p=1.13×10^−10^, ^++^p=0.46, ^+++^p=2.39×10^−12^, and ^#^p=3.90×10^−4^, ^##^p=0.18, ^###^p=1.97×10^−5^. Two independent experiments were performed for each condition. In total, n=20 background measurements, n≥54 single filament measurements, and n≥16 two-filament bundle measurements were taken for each condition. **(C)** Percentage of cells in which Ain1-GFP is observed in actin patches. Two-tailed t-test for data sets with unequal variance yielded ^‡^p-value=0.0029. **(D,E, top)** Fluorescence micrographs of fission yeast in an *ain1*-*1Δ* background overexpressing GFP-tagged wild-type Ain1 **(D)** or mutant Ain1(R216E) **(E)** from the 41Xnmt1 promoter. **(D,E, bottom)** Timelapse (in sec.) of cell end taken from a single Z-plane.

Unlike the in vitro results, the less dynamic Ain1(R216E) mutant was better than wild-type Ain1 at competing with fimbrin Fim1 in vivo. In fission yeast cells expressing endogenous levels of Fim1, overexpressed mutant Ain1(R216E)-GFP localized to actin patches in 100% of cells (Figure 2C,E, Video 2), while overexpressed wild-type Ain1-GFP was localized to actin patches in only ~9% of cells (Figure 2C,D, Video 2). The disparity between Ain1(R216E)’s ability to compete with Fim1 in vitro versus in vivo potentially suggests that slight differences in dynamics may have a bigger effect in a cellular context. In particular, the dynamics of ABPs such as Ain1 may be finely tuned to allow for proper sorting given the large number of actin interacting proteins, with a small change in dynamics skewing the sorting. Alternatively, it is possible that certain aspects of our in vitro system (such as muscle vs fission yeast actin or post-translational modifications of Fim1 (Miao *et al*., 2016)) do not mimic the exact conditions present in vivo.

### Tropomyosin Cdc8 and α-actinin Ain1 do not compete for association with actin filaments

We previously reported that tropomyosin Cdc8, an F-actin side-binding protein that associates with the contractile ring, is displaced from F-actin by fimbrin Fim1 and is therefore prevented from associating with actin patches (Figure 3E-F) (Skau and Kovar, 2010; Christensen *et al*., 2017). However, as fimbrin/plastin and α-actinin isoforms bind to the same site on F-actin, we wondered if Cdc8 and α-actinin Ain1 can associate with actin filaments simultaneously. In multi-color in vitro TIRFM assays, we observed no displacement of Cdc8 from Ain1- or Ain1(R216E)-bundled networks (Figure 3A-D, Video 3), demonstrating that Ain1 and Cdc8 are capable of co-existing on the same F-actin network in vitro, as they do at the contractile ring in cells. Remarkably, TIRFM assays also revealed that Cdc8 actually enhanced the bundling ability of Ain1 (Figure 4G-J). 500 nM Cdc8 increased Ain1-mediated bundling 10-fold over Ain1 alone (Figure 4K, Video 4). This finding is surprising as Ain1 is generally considered to be a poor bundling protein (Li *et al*., 2016), and suggests that the combination of Ain1 and Cdc8 may allow for significant bundling to occur in the context of the contractile ring despite Ain1’s poor bundling ability alone.

**Figure 3:**
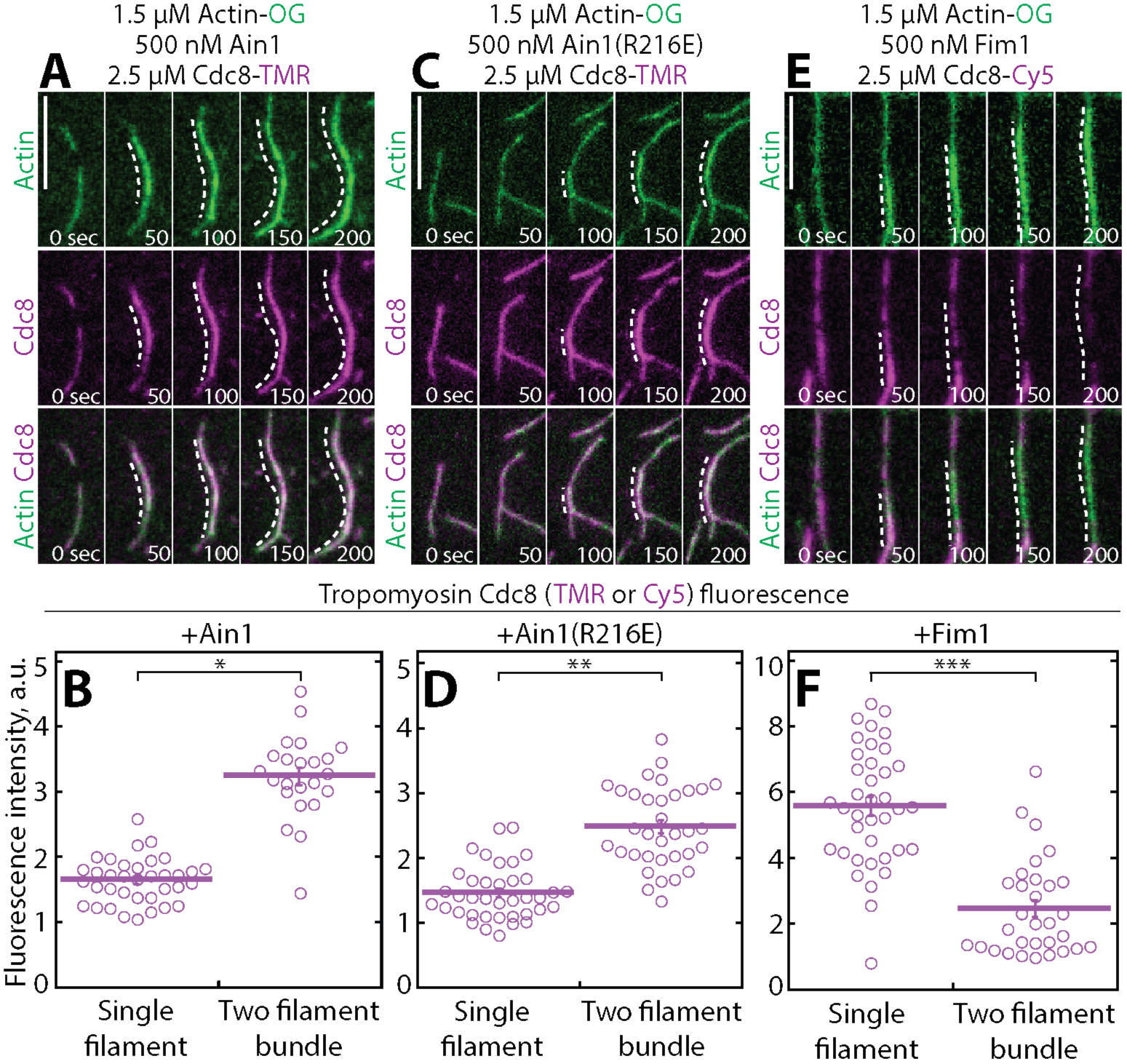
α-actinin Ain1 does not displace Tropomyosin Cdc8 from F-actin bundles *in vitro*. **(A,C,E)** Two-color TIRFM of 1.5 μM Mg-ATP actin (15% Alexa 488-labeled) with 2.5 μM tropomyosin Cdc8 (TMR-labeled) and 500 nM **(A)** wild-type α-actinin Ain1, **(B)** mutant Ain1(R2l6E), or **(C)** fimbrin Fim1 (unlabeled). Scale bars, 1 μm. Dotted lines denote bundled regions. **(B,D,F)** Dot plots of the amount of Cdc8-TMR or Cdc8-Cy5 fluorescence on single filaments or two-filament bundles in the presence of Ain1 **(B)**, Ain1(R216E) **(D),** or Fim1 **(F)** Error bars=s.e. Two-tailed t-tests for data sets with unequal variance yielded p-values ^∗^p=8.24×10^−18^, ^∗∗^p=5.47×10^−12^, ^∗∗∗^p=5.72×10^−11^.

**Figure 4:**
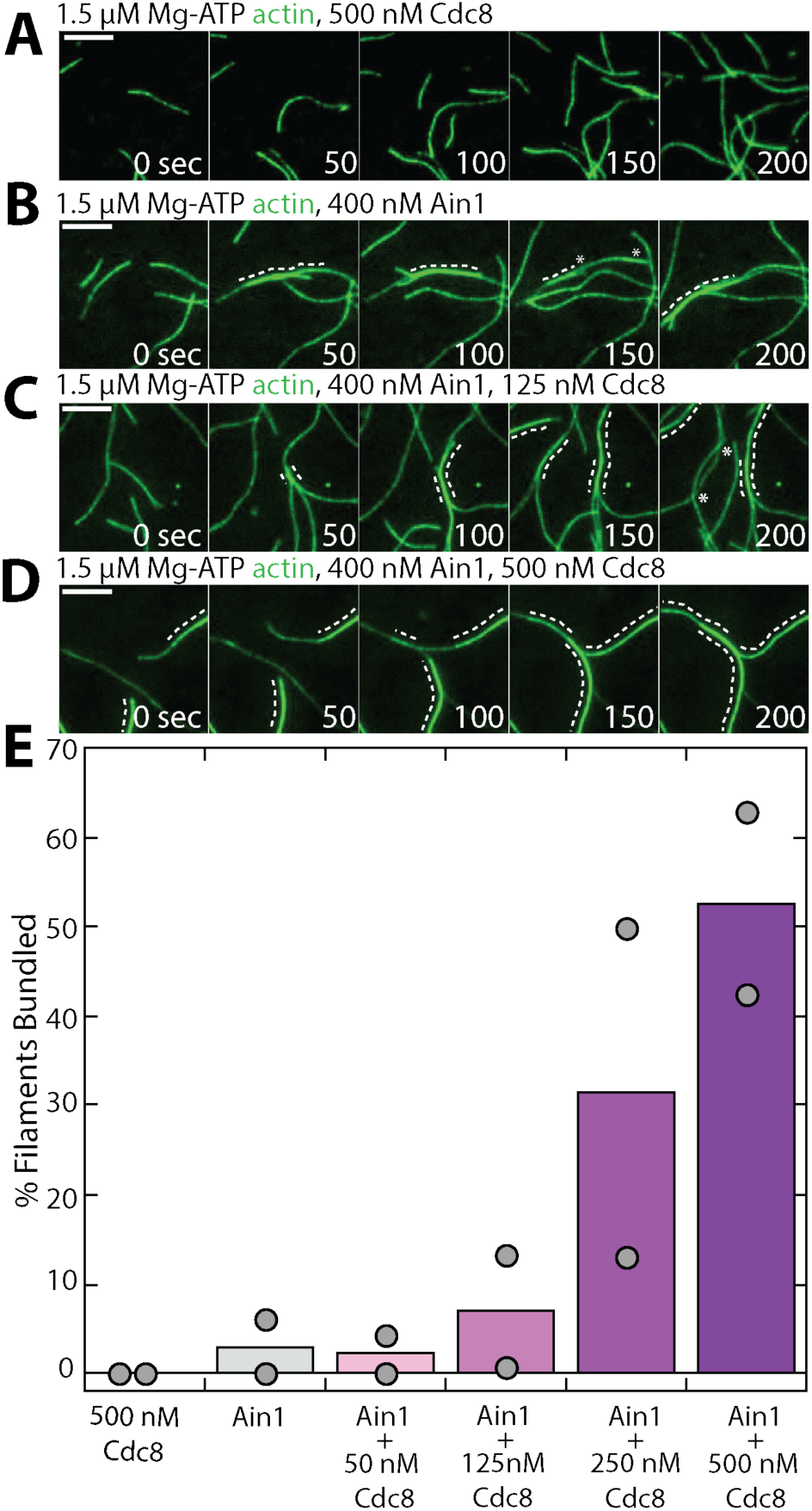
Tropomyosin Cdc8 enhances α-actinin Ain1-mediated F-actin bundling in vitro. (A-D) TIRFM of 1.5 μM Mg-ATP actin (15% Alexa 488-labeled) in the presence of 500 nM Cdc8 **(A)** or 400 nM Ain1 and 0 **(B)**, 125 **(C)**, or 500 nM **(D)** Cdc8. Dotted lines indicate the bundled region. Scale bars, 5 μm. **(E)** Quantification of the percent of bundled F-actin with 400 nM Ain1 and a range of tropomyosin Cdc8 concentrations. Bars indicate averages and gray circles indicate values from independent TIRFM experiments. n=2 independent experiments for each condition.

### Tropomyosin Cdc8 and α-actinin Ain1 cooperate to displace fimbrin Fim1 from actin filaments

On their own, both α-actinin Ain1 and tropomyosin Cdc8 are outcompeted by fimbrin Fim1 for binding to F-actin. Furthermore, there are ~86,500 Fim1 polypeptides in the cell, but only ~3,600 Ain1 molecules (Wu and Pollard, 2005), raising the question as to why more Fim1 is not associated with the contractile ring in wild-type cells. Given that Cdc8 enhances the bundling ability of Ain1, we speculated that the combination of Cdc8 and Ain1 might inhibit Fim1 association with contractile ring actin filaments. We tested this possibility by performing three-color TIRFM with labeled ABPs and quantified Fim1 association with F-actin in the presence of Cdc8 and/or Ain1. As described above, in the absence of other ABPs, Fim1 fully coats F-actin bundles (Figure 2A,B). In the presence of either Cdc8 (Figure 5A,D) or Ain1 (Figure 2A,B) alone, less Fim1 is associated with two-filament bundles, demonstrating that though Fim1 is a better competitor than both Ain1 and Cdc8, both compete to different extents with Fim1 for association with F-actin.

**Figure 5:**
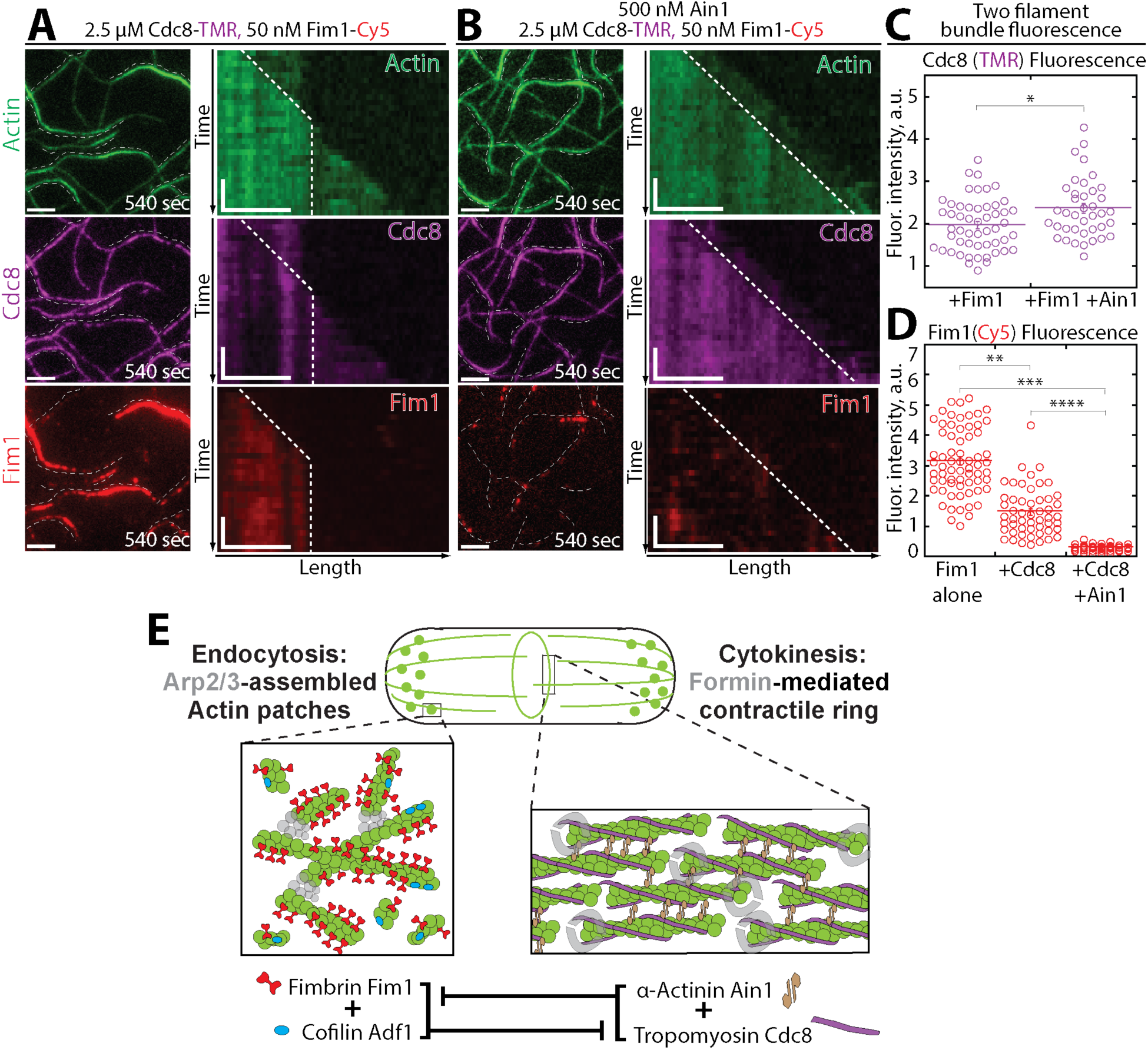
Tropomyosin Cdc8 and α-actinin Ain1 cooperate to compete with fimbrin Fim1 for association with F-actin in vitro. **(A-C)** Three-color TIRFM of 1.5 μM Mg-ATP actin (15% Alexa 488-labeled) with 50 nM fimbrin Fim1 (Cy5-labeled) and 2.5 μM tropomyosin Cdc8 (TMR-labeled) in the **(A)** absence or **(B)** presence of 500 nM unlabeled α-actinin Ain1. **(A-B, left)** Representative TIRF field 540 seconds following reaction initiation. **(A-B, right)** Kymographs of actin, Fim1, and Cdc8 during bundle formation. Dotted lines denote bundled regions. Scale bars, 2.5 μm. Time bar, 30 seconds. **(C-D)** Dot plots of the amount of Cdc8-TMR **(C)** or Fim1-Cy5 fluorescence **(D)** on two-filament bundles in experiments with Cdc8 and Fim1, or Cdc8, Fim1 and Ain1. Purple and red lines denotes mean. Error bars=s.e. Two-tailed t-tests for data sets with unequal variance yielded p-values ^∗^p=0.00716, ^∗∗^p=1.15×10^−15^, ^∗∗∗^p=2.68×10^−30^, ^∗∗∗∗^p=1.14×10^−14^. n≥30 measurements from two independent experiments. **(E)** Model of the involvement of ABP competition in ABP sorting in the fission yeast cell. In endocytic actin patches, fimbrin Fim1 and cofilin Adf1 enhance each other’s activities, resulting in the displacement of tropomyosin Cdc8 from the F-actin network (Christensen *et al*., 2017). In the contractile ring, α-actinin Ain1 and tropomyosin Cdc8 work together to prevent fimbrin Fim1 association with the F-actin network.

In the absence of Ain1, Cdc8 is displaced from F-actin bundles by Fim1 in a cooperative manner, with segments of F-actin bundles completely devoid of Cdc8, concurrent with regions of high Fim1 localization (Figure 5A, Video 5, (Christensen *et al*., 2017)). Conversely, in reactions containing Ain1, ~16% less Cdc8 is displaced by Fim1 from two-filament F-actin bundles (Figure 5B,C, Video 5). Additionally, ~90% less Fim1 is observed to associate with these bundles (Figure 5B,D, Video 5), suggesting that the combination of Ain1 and Cdc8 are capable of preventing Fim1 association with F-actin networks. We speculate that though Ain1 alone is a poor competitor with Fim1, its competition for the same binding site as Fim1 allows it to prevent long stretches of Fim1 from forming that might be capable of displacing Cdc8. The inability of Fim1 to cooperatively associate on F-actin, along with Cdc8’s ability to compete it off of actin filaments result in Fim1 poorly associating with F-actin in the presence of both Cdc8 and Ain1.

## DISCUSSION

### Fimbrin Fim1 association with the contractile ring is regulated by several mechanisms

We have demonstrated that fimbrin Fim1 is prevented from associating with actin filaments by the combined efforts of contractile ring ABPs α-actinin Ain1 and tropomyosin Cdc8. Our model is that Fim1’s association with the contractile ring is inhibited by (1) a preferred association with actin patches, and (2) the combined activity of Cdc8 and Ain1 (Figure 5). If Fim1 preferentially associates with actin patches over other F-actin networks, actin patches could act as a ‘sink’ for Fim1, thereby sequestering Fim1 from associating with other F-actin networks. A preference of Fim1 for actin patches could potentially result from an architectural preference for branched F-actin or a particular twist or conformational change of Arp2/3 complex-assembled F-actin. Alternatively, other upstream ABPs could recruit Fim1 to actin patches by other mechanisms. Future work will involve investigating these upstream signals and their involvement in regulating ABP sorting.

### ABP dynamics mediate competitive interactions

We discovered that the contractile ring ABP α-actinin Ain1 competes with fimbrin Fim1, and that their dynamics regulate their ability associate with different F-actin networks. In particular, we found that a less dynamic mutant Ain1(R216E) localized to F-actin patches in the presence of Fim1. However, in addition to mislocalizing to actin patches, Ain1(R216E)-expressing cells also have cytokinesis defects (Li *et al*., 2016). These findings may explain why fission yeast requires two F-actin bundling proteins. Actin patches require a stable F-actin bundler such as Fim1 to prevent tropomyosin Cdc8 association, and to create boundaries that enhance Adf1-mediated severing (Skau and Kovar, 2010; Christensen *et al*., 2017). On the other hand, a dynamic F-actin bundler is required at the contractile ring to facilitate anti-parallel F-actin contacts while still allowing Cdc8 association and myosin sliding (Li *et al*., 2016). Additionally, binding dynamics seem to mediate Ain1 and Fim1 competition with Cdc8. We demonstrated that Fim1, but not Ain1, displaces Cdc8 from F-actin bundles. However, Fim1 competes with Cdc8 specifically at regions where it binds stably (F-actin bundles), and does not compete as strongly with Cdc8 on single filaments (Christensen *et al*., 2017). Therefore, the presence of a dynamic bundling protein (Fim1 on single filaments and Ain1 on both single and bundled filaments) may allow Cdc8 to remain associated with F-actin in those circumstances.

Additionally, increasing concentrations of Cdc8 actually enhance F-actin bundling in the presence of Ain1 (Figure 3G-K). This bundling enhancement could potentially arise from (1) Cdc8 increasing the residence time of Ain1 on F-actin, or (2) Cdc8 altering the persistence length of F-actin, making the stiffer actin filaments more likely to be incorporated and maintained in a bundle. As we suspect that lowering Ain1 dynamics may negatively affect both contractile ring assembly and constriction (Li *et al*., 2016), we favor the second mechanism.

### Tropomyosin Cdc8 and α-actinin Ain1 work together to compete with fimbrin Fim1 at the contractile ring

Despite the ability of fimbrin Fim1 to actively displace tropomyosin Cdc8 from F-actin bundles (Figure 3E-F), Cdc8 is also capable of inhibiting Fim1 association with F-actin (Figure 5D). Together, we observe that, in our experiments, α-actinin Ain1 and Cdc8 prevent ~90% of Fim1 association with F-actin bundles. Alone, Cdc8 prevents ~50% of Fim1 association while Ain1 alone prevents less than 35% of Fim1 association. It should be noted that though Ain1 and Cdc8 work together to compete with Fim1, our reactions contain low concentrations (50 nM) of Fim1 compared to Ain1 (500 nM or 1 μM) and Cdc8 (2.5 μM). Similarly, given the potent F-actin binding and bundling capabilities of Fim1, a reasonable assumption is that in a cell Cdc8 and Ain1 may only prevent a portion of Fim1 polypeptides from associating with contractile ring F-actin. One possibility is that other ABPs or sets of ABPs at the contractile ring help inhibit Fim1 association. Secondly, budding yeast fimbrin Sac6 is phosphorylated at different stages of the cell cycle, which affects its ability to bundle F-actin (Miao *et al*., 2016). Fission yeast Fim1 might be similarly post-translationally modified, and therefore a portion of the cytoplasmic Fim1 pool might be more or less active. A third non-mutually exclusive possibility is that the actin assembly factors Arp2/3 complex (actin patches) and formin (contractile ring) may bias sorting of ABPs to particular F-actin networks. Future work will seek to determine the contribution of actin assembly factors to ABP sorting, and whether Fim1 and other ABPs are post-translationally modified and how these modifications affect their ability to compete with other ABPs and sort to the correct F-actin network.

## ACKNOWLEDGMENTS

This work was supported by NIH R01 GM079265 and ACS RSG-11-126-01-CSM (to D.R.K.), NIH MCB Training Grant T32 GM0071832 (to J.R.C. and K.E.H.), Initiative for Maximizing Student Development (IMSD) NIGMS R25GM109439 (to M.E.O) and NSF Graduate Student Fellowship DGE-1144082 (to J.R.C.). Additional support was provided to D.R.K. by the University of Chicago MRSEC, funded by the NSF through grant DMR-1420709. We thank Charlie Dulberger and Yujie Li for assistance with Ain1 purification. We also thank Alisha Morganthaler for assistance with preliminary experiments and Jonathan Winkelman, Cristian Suarez, and the Kovar lab for helpful discussions.

## MATERIALS AND METHODS

### Strain construction and growth

Fission yeast strains were created by genetic crossing on SPA5S plates followed by tetrad dissection on YE5S plates. Strains were screened for auxotrophic (leu, ura) or antibiotic (nat, kan) markers and maintained on YE5S plates. Glycerol stocks were created by pelleting cells and resuspending in 750 μL media and 250 μL of 50% sterile glycerol.

### Cell imaging and treatment with CK-666

For live cell imaging, cells were grown in YE5S media overnight at 25°C, subcultured into EMM5S media without thiamine, and kept in log phase for 20-22 hours at 25°C. Cells were imaged directly on glass slides. Z-stacks of 10 slices, 0.5 μm apart were acquired with a 100x, 1.4 NA objective on a Zeiss Axiovert 200M equipped with a Yokogawa CSU-10 spinning-disk unit (McBain, Simi Valley, CA) illuminated with a 50-milliwatt 473-nm DPSS laser, and a Cascade 512B EM-CCD camera (Photometrics, Tucson, AZ) controlled by MetaMorph software (Molecular Devices, Sunnyvale, CA). For CK-666 treatments, CK-666 powder stock (Sigma, St. Louis, MO) was diluted to 10 mM in DMSO. Cells were grown as stated above, and incubated with CK-666 or an equivalent volume of DMSO (control) in a rotator at 25°C for 30 min prior to imaging. Cells were then immediately imaged as above.

### Contractile ring fluorescence quantification

Contractile ring maturation was divided into three stages by measuring the distance between spindle pole bodies (SPBs, visualized by Sad1-tdTomato) and noting constriction of the contractile ring. Stage 1 cells had SPBs less than 6 μm apart, with no observable ring constriction. Stage 2 cells had SPBs greater than 6 μm apart, with no observable ring constriction. Stage 3 cells had SPBs less than 9 μm apart, with evident ring constriction. Quantification of ABP association with defined contractile rings (stages 2 and 3) is shown in Figure 1B-D, and quantification at each distinct ring stage is show in Figure 1, figure supplement 3. The contractile ring region was determined by visually examining the z-stack for the ring site. Normalized contractile ring fluorescence was taken by drawing a region of interest (ROI) around the observed ring and around the whole cell using ImageJ. The mean fluorescence of the ring divided by the whole cell was then determined. A value of 1.00 indicates no increased fluorescence at the site of the contractile ring, while values >1 indicate increased fluorescence at the ring. Maximum projections created in ImageJ (Schindelin *et al*. 2012; Schneider *et al*. 2012) were used for figure images and sum projections were used for quantification.

### Tropomyosin Cdc8 antibody staining

Following standard growth and culturing protocols for live cell imaging, fission yeast cells were stained with anti-Cdc8p (Cranz-Mileva *et al*., 2015). Cells were first fixed in 16% formaldehyde for 5 minutes at 20°C. Cells were then washed in cold 1X PBS and resuspended in 140 μL 1.2M sorbitol. 60 μL fresh protoplasting solution (3 mg/ml zymolase 100T in 1.2M sorbitol) was added and cells were incubated for 7 minutes on a rotator at room temperature. 1 mL of 1% Triton-X was then added to the cells and incubation continued for 2 minutes. Cells were then pelleted and resuspended in 0.5 mL PBAL (10% BSA, 100 mM lysine monohydrochloride, 1 mM NaN_3_, 50 ng/ml ampicillin in PBS) and incubated for 2.5 hours on a rotator at room temperature. Cells were resuspended in 100 μL of anti-Cdc8p 1:10 in PBAL (gift of Sarah Hitchcock-DeGregori) and incubated overnight at 4°C on a rotator. Following incubation with primary antibody, cells were washed 3 times with 0.5 mL PBAL, resuspended in 50 μL Alexa-Flour 555 goat anti-rabbit secondary antibody (Thermo-Fisher Scientific, Carlsbad, CA) (1:100 in PBAL), and incubated for 90 minutes at room temperature on a shaker in the dark. Cells were then washed 5 times with 0.5 mL PBAL and resuspended in 20-30 μL PBAL for imaging. Cells were stored at 4°C and imaged within 48 hours of staining.

### Phallicidin staining

BODIPY-phallicidin staining of fission yeast cells was adapted from Sawin and Nurse, 1998. BODIPY-phallacidin powder (Thermo Fisher Scientific, Waltham, MA) was resuspended to a concentration of 0.2 units/μL in methanol, aliquoted, lyophilized, and stored at −20°C. Fission yeast were grown overnight in YE5S media and fixed in 16% paraformaldehyde for 5 minutes at room temperature. Cells were washed with room temperature PEM buffer 3 times and permeabilized in PEM with 1% triton X-100 for exactly 1 minute. Cells were then spun for 30 seconds at 7000 RPM. Cells were washed in PEM buffer 3 times and resuspended in 10 μL PEM buffer. Lyophilized BODIPY-phallacidin was resuspended to 1 unit/μL. 1 μL (1 unit) of resuspended phallacidin was added to 10 μL of cells in PEM buffer and incubated in the dark for 30 minutes at room temperature. Following incubation in the dark, cells were washed with 1 mL PEM and spun at 7000 RPM for 30 seconds. Supernatant was removed and cells resuspended in a small volume. For cells stained with anti-Cdc8p and BODIPY-phallicidin, cells were first treated with primary and secondary antibodies, washed with PBAL, and then stained with BODIPY-phallicidin.

### Protein purification

Chicken skeletal muscle actin was purified as described previously (Spudich and Watt, 1971). Fimbrin Fim1 and tropomyosin AlaSer-Cdc8 (WT and I76C mutant) were expressed in BL21-Codon Plus (DE3)-RP (Agilent Technologies, Santa Clara, CA). His-tagged Fim1 was purified using Talon Metal Affinity Resin (Clontech, Mountain View, CA) (Skau and Kovar, 2010). Cdc8 was purified by boiling the cell lysate, performing an ammonium sulfate cut, and running on an anion exchange column (Skau and Kovar, 2010). His-tagged wild-type α-actinin Ain1 and mutant Ain1(R216E) were expressed in High Five insect cells using baculovirus expression and purified using Talon Metal Affinity Resin (Li *et al*., 2016).

The A_28_O of purified proteins was taken with a Nanodrop 2000c Spectrophotometer (Thermo-Scientific, Waltham, MA). Protein concentration was calculated using extinction coefficients Fim1: 55,140 M^−1^ cm^−1^, Cdc8 (WT and I76C mutant): 2,980 M^−1^ cm^−1^, Ain1 and Ain1(R216E): 86477 M^−1^ cm^−1^. Proteins were labeled with TMR-6-maleimide (Life Technologies, Grand Island, NY) or Cy5-monomaleimide (GE Healthcare, Little Chalfont, UK) dyes following manufacturer’s protocols following purification. Proteins were flash-frozen in liquid nitrogen and stored at −80°C.

### TIRF microscopy

Time-lapse TIRFM movies were obtained using an Olympus IX-71 microscope with through-the-objective TIRF illumination, iXon EMCCD camera (Andor Technology), and a cellTIRF 4Line system (Olympus). The actin binding proteins (ABPs) of interest were initially added to a polymerization mix (10 mM imidazole (pH 7.0), 50 mM KCl, 1 mM MgCl_2_, 1 mM EGTA, 50 mM DTT, 0.2 mM ATP, 50 μM CaCl_2_, 15 mM glucose, 20 μg/mL catalase, 100 μg/mL glucose oxidase, and 0.5% (400 centipoise) methylcellulose). This ABP/polymerization mix was then added to Mg-ATP-actin (15% Alexa 488-labeled) to induce F-actin assembly in the presence of the ABPs of interest. The mixture was then added to a flow chamber and imaged at room temperature at 5 s intervals (unless otherwise noted).

### Quantification of bundling

The percentage of actin filaments bundled was quantified at similar actin filament densities (between 2095 and 2295 μm total filament length) for each experiment. The total actin filament length in the chamber was measured manually by creating ROIs for every actin filament and measuring total actin filament length in FIJI (Schindelin *et al*., 2012; Schneider *et al*., 2012). ROIs for every segment of actin filament present in a bundle were then created and total bundled filament length measured. The ratio of actin filament present in a bundle vs. total actin filament length was then calculated.

### Quantification of fluorescence intensity on actin filaments or bundles

Fluorescence intensity on actin filaments was quantified on movies taken under the same microscope conditions (laser intensity and angle, exposure time) and with the same protein batches. Fluorescence intensity was quantified at the same time point in each compared movie. The actin channel was used to identify single actin filaments or two-filament actin bundles, and ROIs of a 3-pixel segmented line were created along all single filaments or two-filament bundles in the selected frame. The mean fluorescence for each segment was then measured using ImageJ.

### Quantifying number of cells with Ain1 in actin patches

To quantify the number of cells containing Ain1-GFP in actin patches, one minute timelapse movies of 1 frame per second were taken, imaging both Ain1-GFP and an actin patch marker (ArpC5-mCherry or Fim1-mCherry). Movie files for independent experiments and replicates were blinded and independently analyzed for number of cells containing Ain1-GFP in actin patches using FIJI (Schindelin *et al*., 2012; Schneider *et al*., 2012). For a single cell to count as positively containing Ain1-GFP in actin patches, three criteria had to be met: 1) at least one distinguishable actin patch containing Ain1-GFP was observed, 2) the observed actin patch(es) contained Ain1-GFP for at least 3 frames and 3) the Ain1-GFP signal trajectory matched the channel expressing either ArpC5-mCherry or Fim1-mCherry. Total number of cells and cells with actin patches containing Ain1-GFP were then calculated to obtain percent of cells containing Ain1-GFP in actin patches.

**Figure 1 - figure supplement 1.**
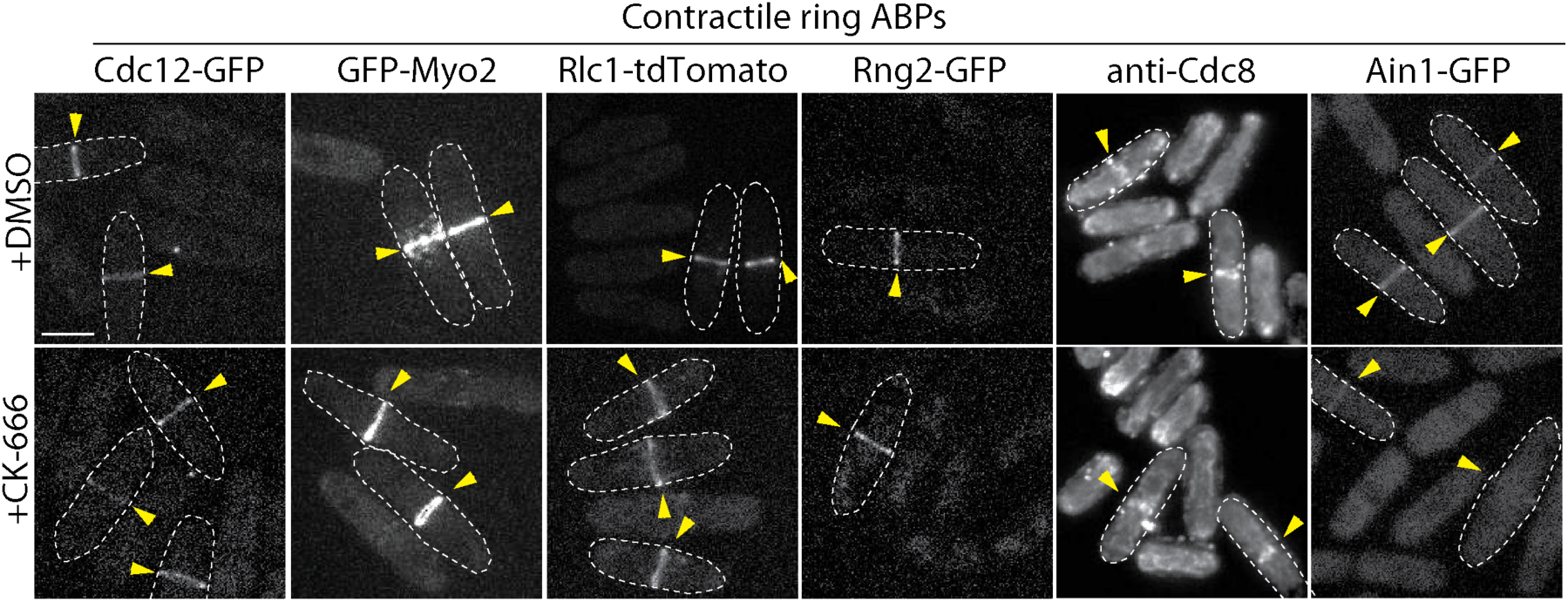
Contractile ring ABP localization following CK-666 treatment. Fluorescent micrographs of fission yeast cells either immuno-stained (anti-Cdc8) or expressing fluorescently-tagged ABPs from the endogenous locus. Cells were treated with DMSO (control) or 200 μM Arp2/3 complex inhibitor CK-666. Yellow arrowheads denote contractile rings. Dotted lines outline individual cells for clarity.

**Figure 1 - figure supplement 2.**
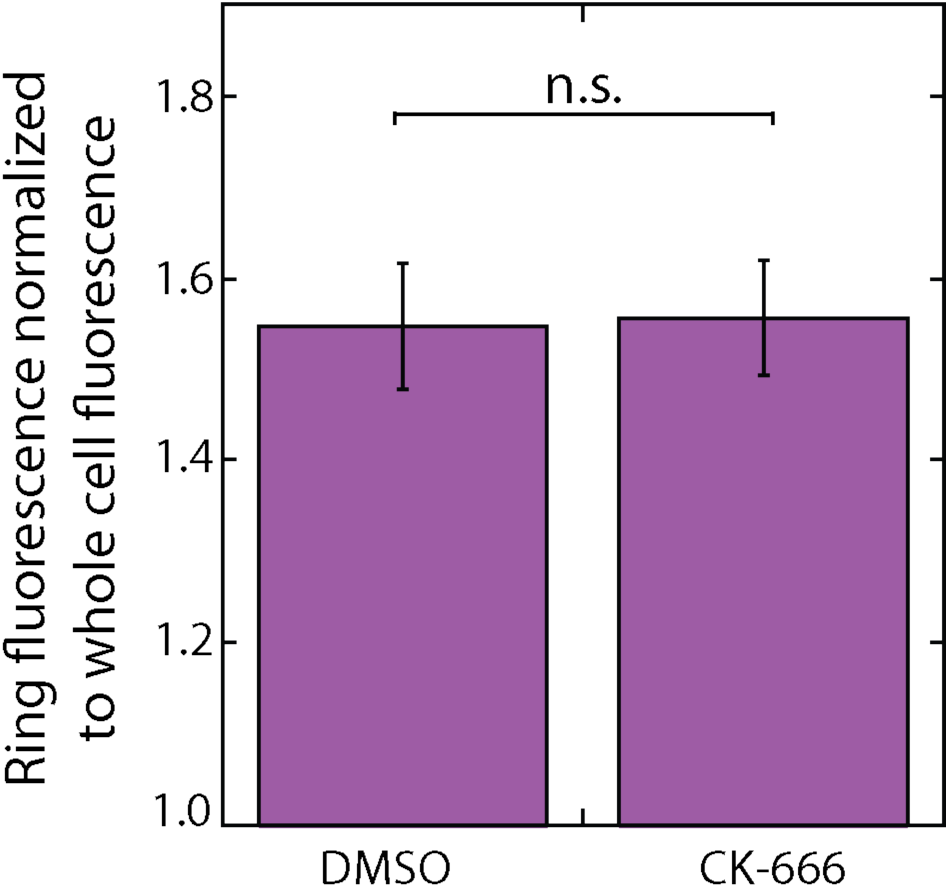
Tropomyosin Cdc8 does not leave the contractile ring following CK-666 treatment. Mean anti-Cdc8 contractile ring fluorescence normalized to total cell fluorescence. Error bars=s.e. Two-tailed t-test for data sets with unequal variance yielded p-value=0.9338. n≥10 cells in each condition.

**Figure 1 - figure supplement 3.**
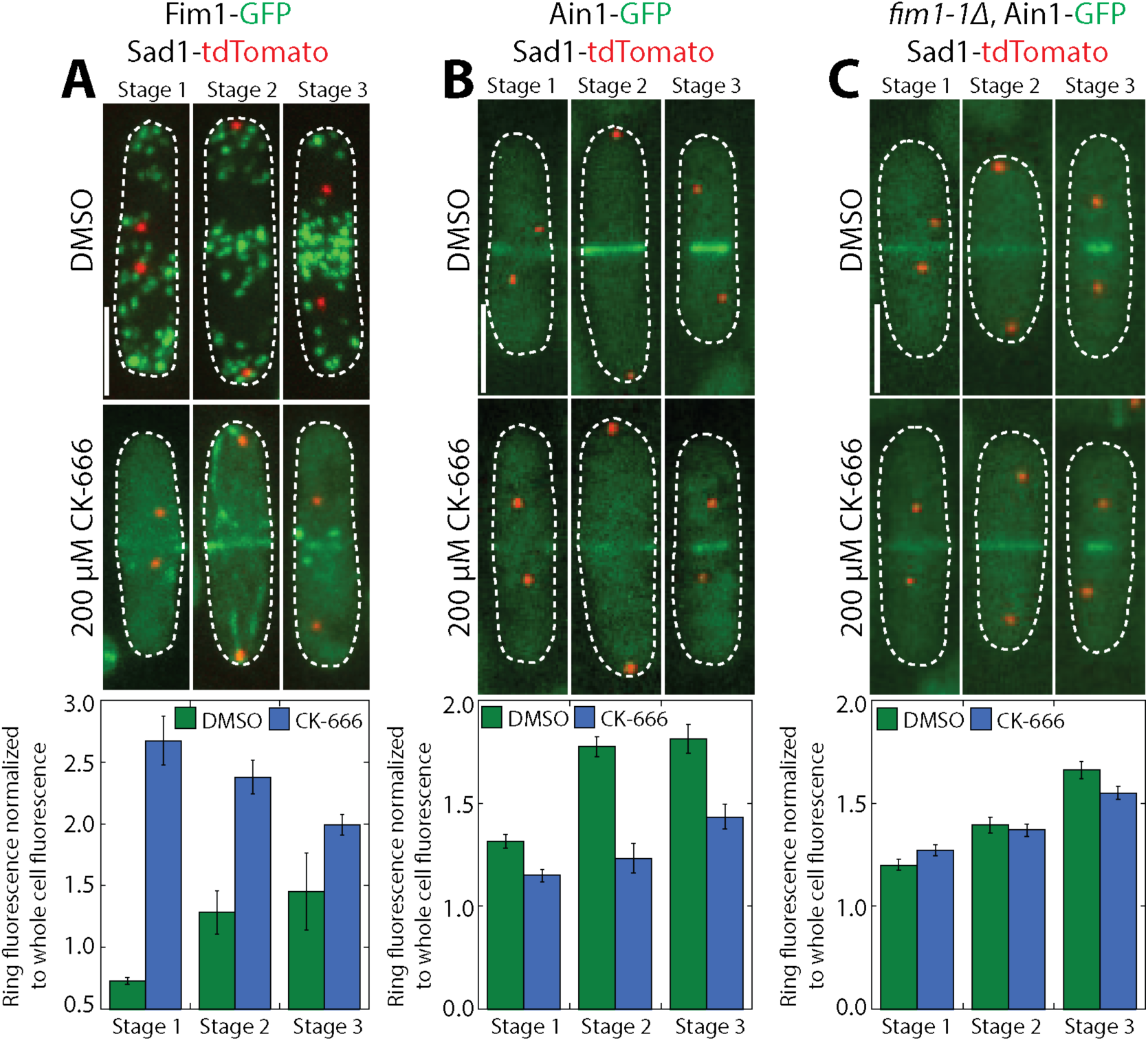
Fimbrin Fim1 displaces α-actinin Ain1 from the contractile ring following CK-666 treatment. **(A-C, top)** Fluorescent micrographs of fission yeast cells expressing spindle pole body marker Sad1-tdTomato and Fim1-GFP **(A)**, Ain1-GFP **(B)**, or Ain1-GFP in a *fim1*-*1Δ* background **(C),** following treatment with DMSO (control, top) or 200 μM Arp2/3 complex inhibitor CK-666 (bottom). Scale bars, 5 μm. **(A-C, bottom)** Mean Fim1-GFP **(A)** or Ain1-GFP **(B,C)** contractile ring fluorescence normalized to whole cell fluorescence for cells in stage 1 (contractile ring formation), stage 2 (contractile ring dwell), or stage 3 (contractile ring constriction) of cytokinesis following treatment with DMSO (control) or 200 μM CK-666. Quantification of cells from stages 2 and 3 are also shown in Figure 1B-D. Dotted lines outline cells. Error bars=s.e. n≥6 cells for each condition.

**Table 1:** Fission yeast strains used in this study.

| Strain name | Genotype | Reference |
| --- | --- | --- |
| KV91 | h+, kanMX6::myo2p::gfp-myo2p+, ade6-M210, leu1-32, ura4-D18 | Wu *et al.*, 2003 |
| KV343 | h?, cdc12-mGFP::KanR | This study |
| KV459 | h+, rlc1-tdTomato-natMX6 ade6-M210 leu1-32 ura4-D18 | This study |
| KV579 | h-, ain1-GFP-KanMX6, URA+ | Wu *et al.*, 2001 |
| KV588 | h+, pAct1 Lifeact-GFP::Leu+; ade6-m216; leu1-32; ura4-D18 | Huang *et al.*, 2012 |
| KV707 | h-, leu1-32, his3-D1, ura4-D18, ade6-M216, Pnmt41-SpAin1-mGFP::Leu+ | This study |
| KV804 | h?, fim1-mCherry-natMX6, ain1-Δ1:: kanMX6, Pnmt41-SpAin1-mGFP::Leu+ | This study |
| KV818 | h+ kanMX6-Prng2-mEGFP-rng2 ade6-M210 leu1-32 ura4-D18 | Laporte *et al.*, 2011 |
| KV856 | h?, ain1-Δ1:: kanMX6, Pnmt41-SpAin1(R216E)-mGFP::Leu+, fim1-mCherry-natMX6, ade6, leu1-32, ura4-D18 | This study |
| KV861 | h?, ain1-GFP-kanMX6, sad1-tdTomato-natMX6, ade6-m21?, leu1-32, ura4-D18 | This study |
| KV878 | h+, fim1-mGFP-kanMX6, sad1-tdTomato-natMX6 | This study |
| KV908 | h? fim1-1Δ-kanMX6, ain1-GFP-kanMX6, sad1-tdTomato::natMX6 | This study |
| KV963 | h?, fim1-1Δ::kanMX6, Pnmt41-SpAin1-mGFP::Leu+ | This study |
| KV968 | h? Pnmt41-SpAin1-mGFP::Leu+, ARPC5-mCherry-natMX6 | This study |

## Video Legends

### Video 1. Overexpressed Ain1-GFP localizes to actin patches in the absence of fimbrin, related to Figure 1.

Fission yeast cells expressing actin patch marker ArpC5-mCherry (right) with Ain1-GFP (left) at endogenous or overexpressed levels. (Top panels) Endogenous Ain1-GFP in a *fim1-1Δ* background. (Middle panels) Ain1-GFP overexpressed under the 41xnmt promoter in a *fim1-1Δ* background. (Bottom panels) Ain1-GFP overexpressed under the 41xnmt promoter in a wild-type background (bottom). Yellow arrowheads indicate cell ends with Ain1-GFP in actin patches. Scale bar, 5 μm. Time in seconds.

### Video 2. Overexpressed mutant Ain1(R216E)-GFP, but not Ain1-GFP, localizes to actin patches in the presence of fimbrin, related to Figure 2

(Top) Fission yeast cells with Ain1-GFP overexpressed under the 41xnmt promoter (left) and expressing Fim1-mCherry at the endogenous locus (right). (Bottom) Fission yeast cells with Ain1(R216E)-GFP overexpressed under the 41xnmt promoter and expressing Fim1-mCherry at the endogenous locus. Scale bar, 5 μm. Time in seconds.

### Video 3. α-actinin Ain1 does not displace tropomyosin Cdc8 from F-actin bundles, related to Figure 3

TIRFM of 1.5 μM actin (Alexa-488 labeled) (left column) with 2.5 μM tropomyosin Cdc8 (TMR- or Cy5-labeled) (middle column) and 500 nM Ain1 (top row), 500 nM Ain1(R216E) (middle row), or 500 nM Fim1 (bottom row). Scale bar, 5 μm. Time in sec.

### Video 4. Tropomyosin Cdc8 enhances α-actinin Ain1-mediated bundling, related to Figure 4

TlRFM of 1.5 μM actin (Alexa-488 labeled) with varying concentrations of unlabeled Ain1 and Cdc8. Scale bar, 5 μm. Time in sec.

### Video 5. Tropomyosin Cdc8 and α-actinin Ain1 work together to prevent fimbrin Fim1 from associating with F-actin bundles, related to Figure 5

Three-color TIRFM of 1.5 μM actin (Alexa-488 labeled), 2.5 μM tropomyosin Cdc8 (Cy5-labeled) and 50 nM fimbrin Fim1 (TMR-labeled) with (bottom) or without (top) 500 nM Ain1 (unlabeled). Scale bar, 5 μm. Time in sec.

